# Full-length 16S profiling reveals individualized gut microbiota dynamics during short-duration spaceflight

**DOI:** 10.64898/2026.09.01.748461

**Authors:** Gábor Gulyás, Balázs Kakuk, Ákos Dörmő, István Prazsák, Tamás Járay, Zsolt Csabai, Ádám Tibor Schlégl, Zsolt Boldogkői, Dóra Tombácz

## Abstract

Human spaceflight may perturb the gut microbiota, but densely sampled short missions remain poorly characterized. We profiled 27 phase-matched fecal samples from two astronauts during an 18-day International Space Station mission and one ground-based participant following the same daily schedule using Oxford Nanopore full-length 16S sequencing. Participant identity dominated genus-level Bray-Curtis variation (R2 = 0.489, p < 0.001). In astronaut-only community analyses, mission phase explained 23.4% of genus-level (p = 0.035) and 22.3% of species-level (p = 0.021) variation. Astronauts showed greater displacement from personal baselines than B1 (0.331 versus 0.171) and 1.58-fold higher volatility. Astronaut-only taxon models identified 2 of 81 genera and 5 of 139 species; *Collinsella* increased from quarantine to orbit (coefficient = 2.586, q = 0.037). Thus, the short-duration spaceflight interval was accompanied by individualized, temporally localized community and taxon shifts rather than uniform microbiota restructuring.

## Introduction

Human spaceflight is entering a period of increasing mission complexity and broader participation in orbital flight. Understanding biological responses relevant to crew health is therefore a central objective of space medicine. Recurring features of spaceflight include immune dysregulation, oxidative stress, mitochondrial and epigenetic alterations, telomere dynamics and microbiome shifts.1,2

The gut microbiota contributes to nutrient metabolism, immune regulation, epithelial barrier maintenance and colonization resistance. Perturbations of this ecosystem may influence adaptation, infection susceptibility and recovery after flight. However, gut communities are also highly individualized, and mission-associated changes must be interpreted against persistent personal microbial backgrounds.3,4,5

The spaceflight environment combines multiple potential drivers of microbiota change, including microgravity, elevated radiation exposure, confinement, altered diet, circadian disruption, psychological stress and reduced environmental microbial exposure. These factors may affect microbial communities directly or indirectly through host physiology, and their individual contributions are difficult to separate in astronaut studies.6,7,8

Previous human spaceflight studies have reported transient or persistent changes in host-associated microbial communities, while consistently emphasizing substantial inter-individual variation. The NASA Twins Study and longitudinal ISS investigations identified mission-associated gut and multi-site microbiome changes, and the Inspiration4 study demonstrated that measurable host and microbiome responses can occur during short-duration flight.9,10,11,12

Ground-based analog studies provide complementary evidence that confinement and restricted environmental exposure can reshape gut microbial composition even without microgravity. In the Mars500 project, gradual community changes developed during prolonged isolation while participant-specific characteristics remained detectable. Together, these studies indicate that astronaut microbiome responses should be assessed as individualized longitudinal trajectories rather than uniform group-level shifts.13

Full-length 16S rRNA gene sequencing is well suited to operationally constrained longitudinal monitoring. V1-V9 amplicons can provide improved taxonomic resolution over short hypervariable-region assays while requiring less input material and sequencing depth than shotgun metagenomics. Nanopore sequencing has also been used for culture-independent microbial monitoring aboard the International Space Station, and workflow comparisons show that extraction, sequencing and bioinformatic choices substantially shape reported taxonomic profiles.14,15,16

Here, we used Oxford Nanopore full-length 16S profiling to characterize fecal microbiota dynamics in two astronauts participating in a 20-day spaceflight mission (20 days and 3 hours in total, including 2 days and 6 hours of free flight and 17 days and 21 hours aboard the International Space Station) and one ground-based participant sampled across corresponding time windows. We evaluated global community structure, displacement from individual preflight baselines, longitudinal volatility, and baseline-adjusted taxon trajectories. Primary inference was restricted to the predefined preflight, in-flight, and early postflight windows, while later travel-associated samples were reserved for sensitivity analysis.

## Results

### Study design and primary analysis framework

The study included two astronauts who completed a 20-day spaceflight mission (20 days and 3 hours in total), including 17 days and 21 hours aboard the International Space Station, and one ground-based participant. (Fig. 1) The complete sequencing dataset comprised 47 samples. For primary fecal microbiome inference, we retained 27 phase-matched samples representing five preflight quarantine windows (Q1-Q5), one in-flight window (O1) and three early postflight recovery windows (PostF1-PostF3; Table 1; Supplementary Table 1). PostF4-PostF6 were excluded from primary inference because they coincided with substantial post-mission travel and were treated as a potentially travel-confounded late-recovery interval.

**Figure 1.**
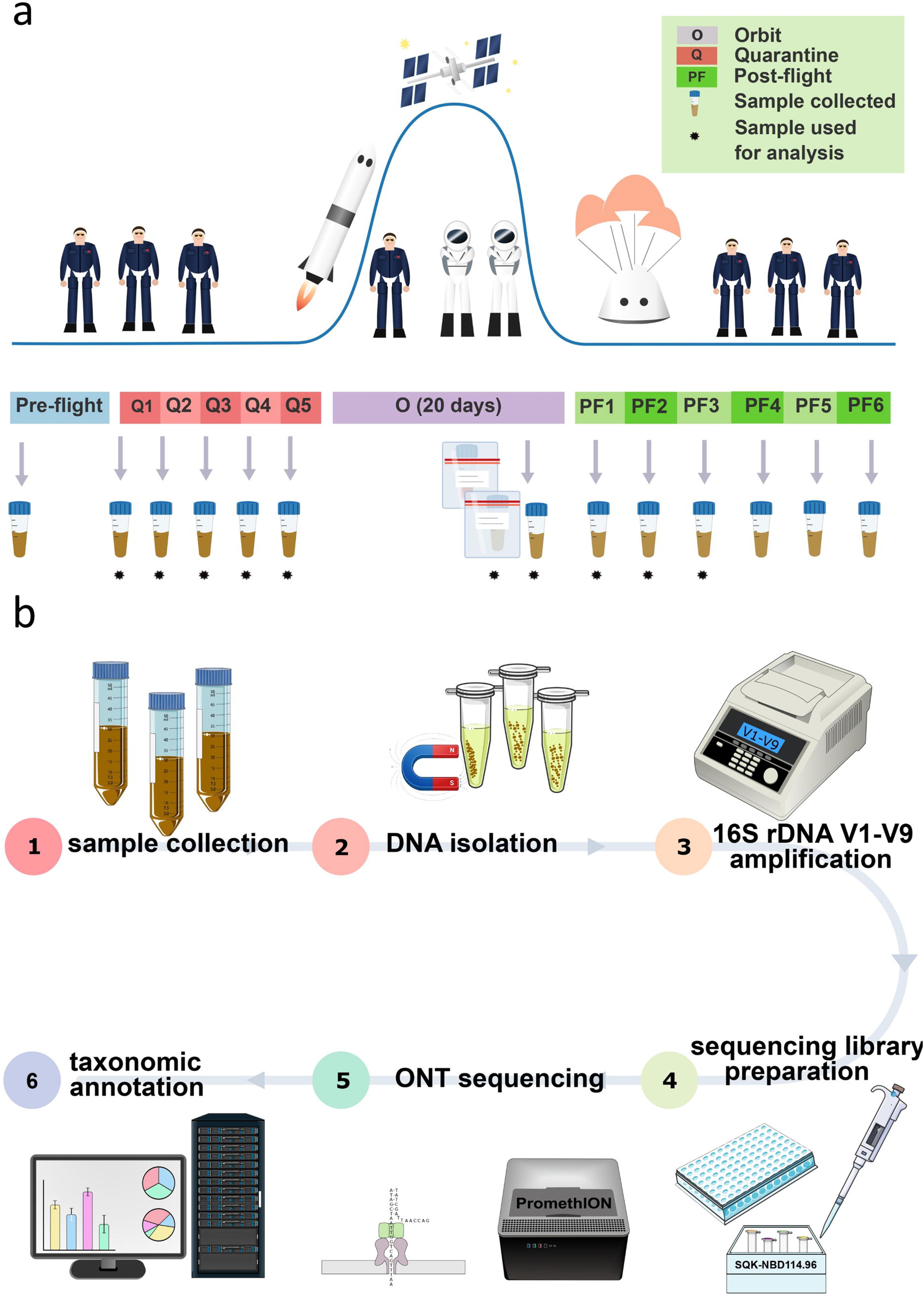
Study design and analytical workflow. Overview of the study design, longitudinal sampling scheme, laboratory workflow and bioinformatic analysis pipeline.

**Table 1.**
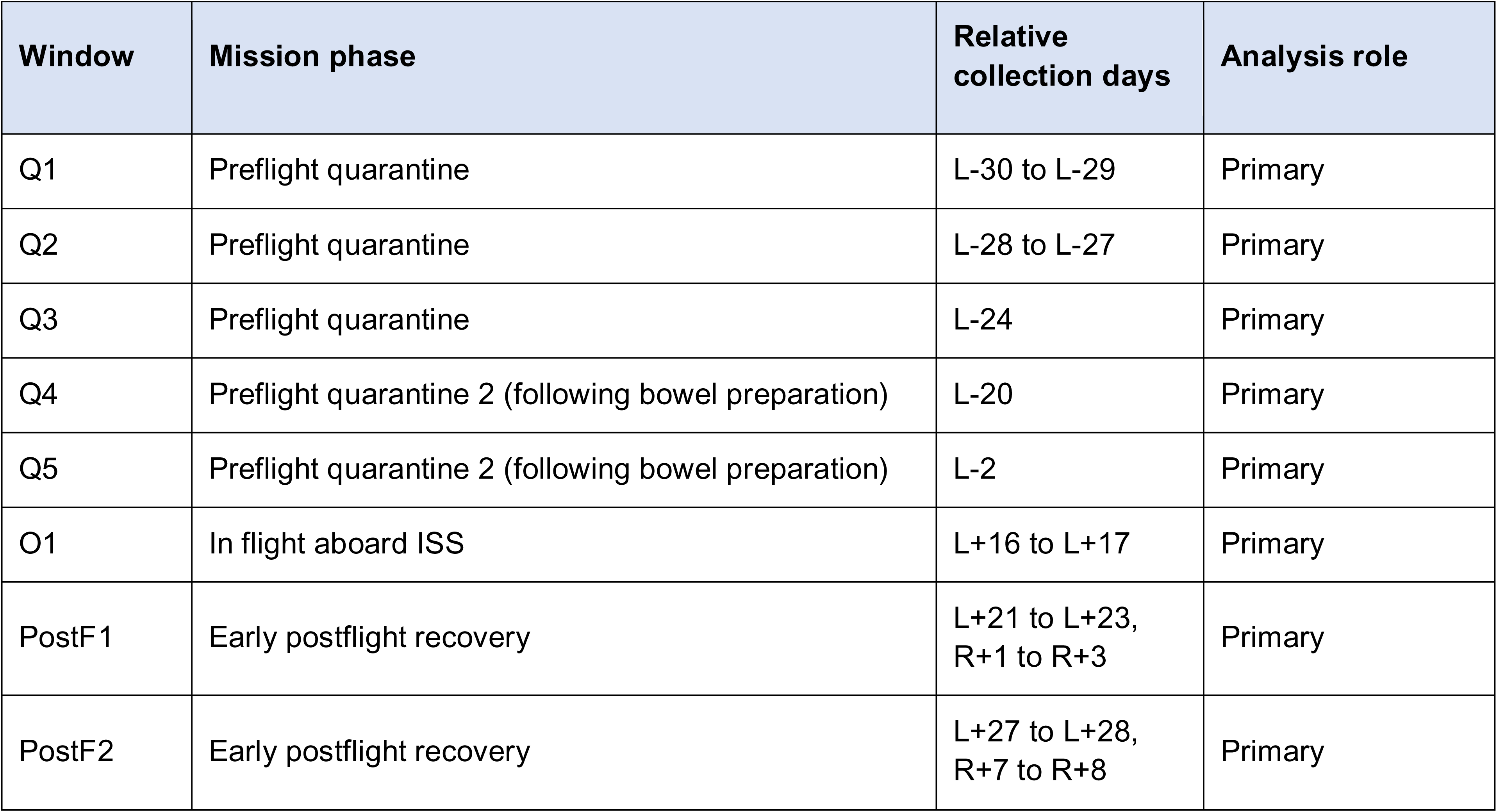
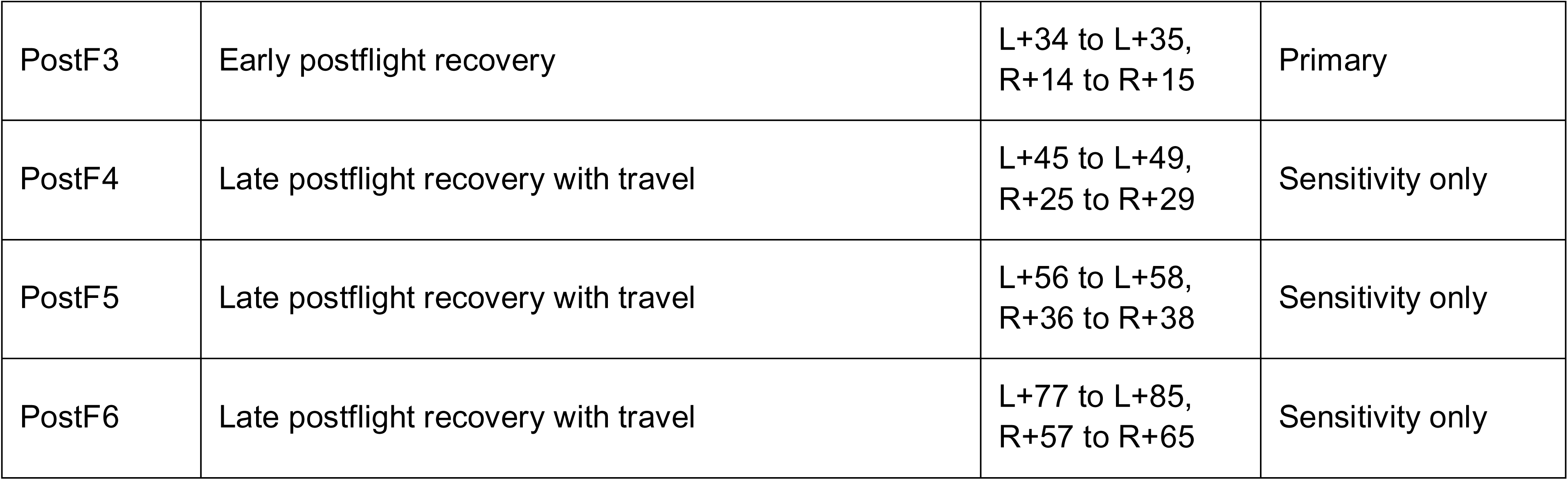
Phase-matched fecal sampling windows and analysis role.

| Window | Mission phase | Relative collection days | Analysis role |
| --- | --- | --- | --- |
| Q1 | Preflight quarantine | L-30 to L-29 | Primary |
| Q2 | Preflight quarantine | L-28 to L-27 | Primary |
| Q3 | Preflight quarantine | L-24 | Primary |
| Q4 | Preflight quarantine 2 (following bowel preparation) | L-20 | Primary |
| Q5 | Preflight quarantine 2 (following bowel preparation) | L-2 | Primary |
| O1 | In flight aboard ISS | L+16 to L+17 | Primary |
| PostF1 | Early postflight recovery | L+21 to L+23,<br>R+1 to R+3 | Primary |
| PostF2 | Early postflight recovery | L+27 to L+28,<br>R+7 to R+8 | Primary |
| PostF3 | Early postflight recovery | L+34 to L+35,<br>R+14 to R+15 | Primary |
| PostF4 | Late postflight recovery with travel | L+45 to L+49,<br>R+25 to R+29 | Sensitivity only |
| PostF5 | Late postflight recovery with travel | L+56 to L+58,<br>R+36 to R+38 | Sensitivity only |
| PostF6 | Late postflight recovery with travel | L+77 to L+85,<br>R+57 to R+65 | Sensitivity only |

PromethION sequencing of the complete dataset generated 64.98 million reads, of which 60.3 million passed sequencing-level quality filtering. After sample selection and the Kraken2 analysis filters described below, the primary 27-sample phyloseq object contained 23,820,971 reads (mean 882,258; median 930,952; range 208,840-2,039,644 reads per sample).

### Broad phylum-level composition remained dominated by the same major taxa across mission phases

Phylum-level relative abundance profiles showed no major restructuring of the broad taxonomic composition across the four primary mission phases (Fig. 2). *Bacillota* remained the dominant phylum in all three participants, with *Bacteroidota* representing the second largest component and *Actinomycetota* contributing a smaller but variable fraction. The relative abundances of these major phyla fluctuated across Quarantine 1, Quarantine 2, Orbit and Postflight 1, with participant-specific differences in the magnitude and direction of these changes. Such within-individual temporal fluctuations are well documented in longitudinal human gut microbiome studies,^17,18^ and no phase was characterized by a large-scale replacement of the dominant phylum-level community structure. Overall, the same dominant phyla were retained across mission phases despite participant-specific fluctuations in their relative abundances, with no wholesale replacement of the broad phylum-level community structure.

**Figure 2.**
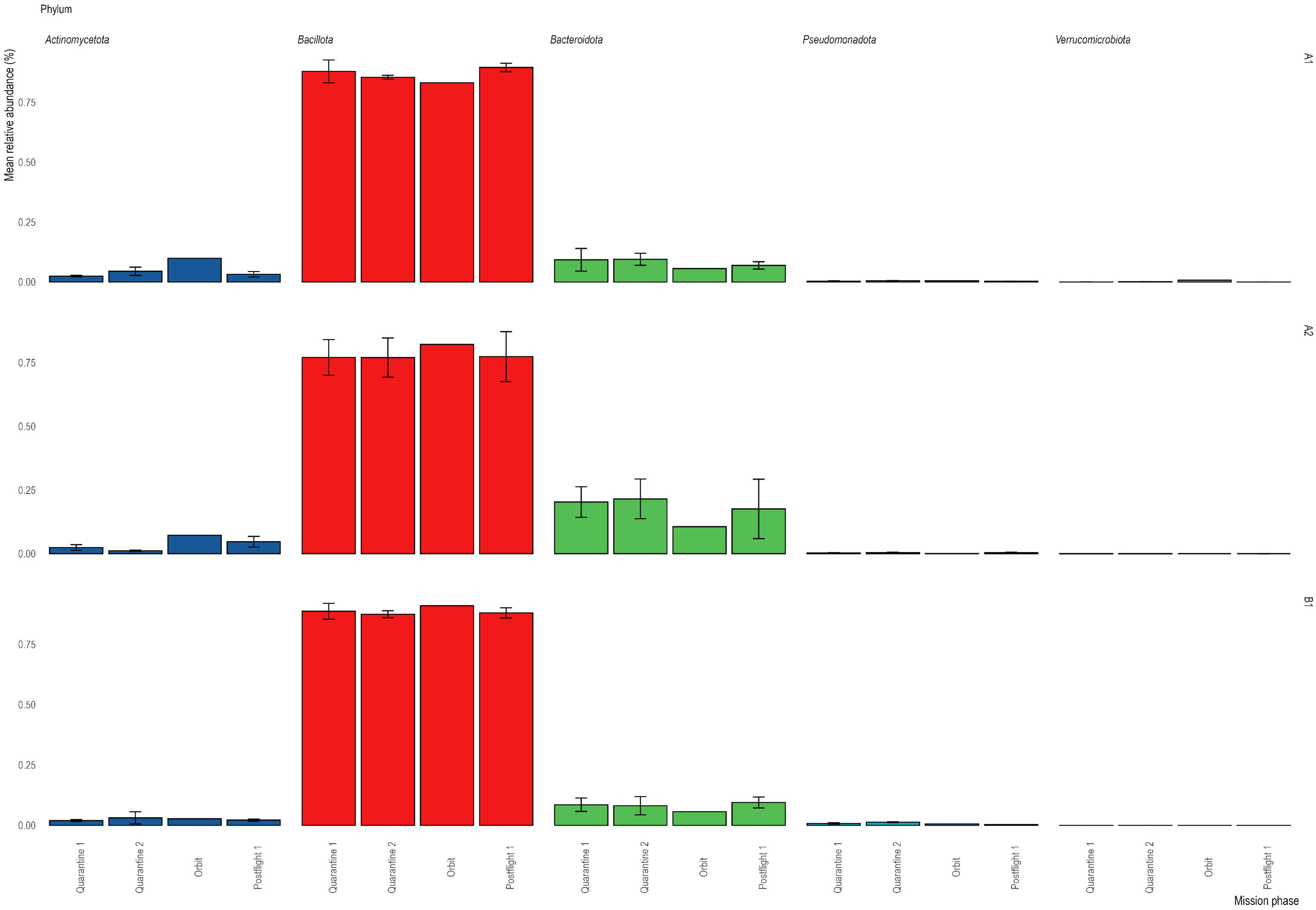
Phylum-level composition across mission phases. Mean relative abundances of the dominant phyla are shown for astronauts A1 and A2 and the ground-based participant B1 across Quarantine 1, Quarantine 2, Orbit and Postflight 1. Phase means were calculated from three sampling windows in Quarantine 1, two in Quarantine 2, one in Orbit and three in Postflight 1 for each participant. Error bars indicate the standard deviation across sampling windows where more than one window was available.

### Participant identity dominated beta diversity

Genus-level Bray-Curtis NMDS of the 27-sample A1/A2/B1 dataset separated samples mainly by participant and yielded a stress value of 0.098 (Fig. 3). B1 followed the most compact longitudinal trajectory, whereas A1 and A2 showed broader movement in ordination space around the in-flight and early-recovery windows. In the full three-participant marginal PERMANOVA, participant identity explained 48.9% of genus-level variation (p = 0.0001), whereas mission phase explained 9.6% (p = 0.0758). Because the manuscript-facing phase question concerns the flown participants, we used the A1/A2 astronaut-only PERMANOVA as the primary community-phase test. In this astronaut-only analysis, mission phase explained 23.4% of genus-level variation (p = 0.035) and 22.3% of species-level variation (p = 0.021), with similar but borderline effects at phylum and order levels (Table 2). Thus, overall community structure remained individualized, but the astronauts showed a detectable mission-phase-associated community shift at higher taxonomic resolution. Astronaut-only mission-phase PERMDISP was non-significant at genus (p = 0.7725) and species (p = 0.3215) levels. Full-rank PERMANOVA and PERMDISP details are provided in Supplementary Tables 2-4, with Aitchison PCA and alpha-diversity sensitivity summaries in Supplementary Figures 1 and 2.

**Figure 3.**
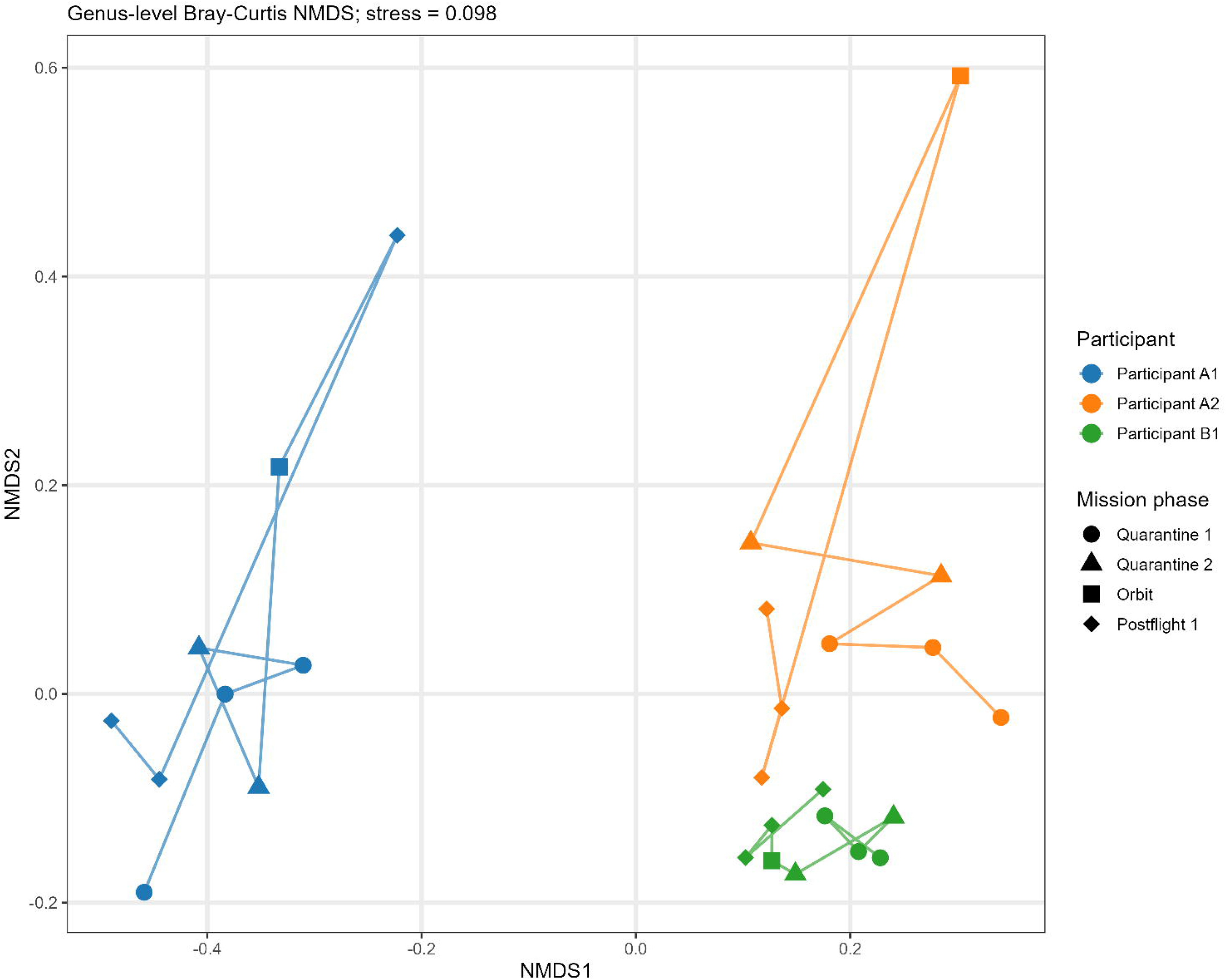
Genus-level Bray-Curtis NMDS trajectories. Each point represents one primary fecal sample, lines connect successive windows within a participant and point shape indicates mission phase. The two-dimensional solution had a stress value of 0.098. The figure displays all 27 samples retained in the primary analysis (Q1–PostF3).

**Table 2.** Community-level PERMANOVA and mission-phase dispersion across taxonomic ranks.

| Rank | Full cohort: Participant R<sup>2</sup> (p) | Full cohort: Phase R<sup>2</sup> (p) | Astronaut-only: Phase R<sup>2</sup> (p) | Astronaut-only: PERMDISP p |
| --- | --- | --- | --- | --- |
| Phylum | R<sup>2</sup> = 0.483;<br>p = 0.0003 | R<sup>2</sup> = 0.067;<br>p = 0.3781 | R<sup>2</sup> = 0.327;<br>p = 0.057 | p = 0.8637 |
| Order | R<sup>2</sup> = 0.433;<br>p = 0.0001 | R<sup>2</sup> = 0.091;<br>p = 0.2101 | R<sup>2</sup> = 0.312;<br>p = 0.052 | p = 0.9322 |
| Genus | R<sup>2</sup> = 0.489;<br>p = 0.0001 | R<sup>2</sup> = 0.096;<br>p = 0.0758 | R<sup>2</sup> = 0.234;<br>p = 0.035 | p = 0.7725 |
| Species | R<sup>2</sup> = 0.541;<br>p = 0.0001 | R<sup>2</sup> = 0.087;<br>p = 0.0780 | R<sup>2</sup> = 0.223;<br>p = 0.021 | p = 0.3215 |

### Astronauts showed greater displacement from personal baseline and higher volatility

Each participant was compared with the centroid of their own Q1-Q5 genus profile. At O1, the mean Bray-Curtis distance of the two astronauts from their personal baselines was 0.331, compared with 0.171 in B1 (Fig. 4a). The corresponding astronaut mean remained elevated at PostF1 (0.330 versus 0.163), approached the B1 value at PostF2 (0.190 versus 0.196) and was again higher at PostF3 (0.213 versus 0.120). These comparisons are descriptive because they average two flown participants against one ground-based participant. The Bray-Curtis and Aitchison baseline-distance summaries are provided in Supplementary Table 5.

**Figure 4.**
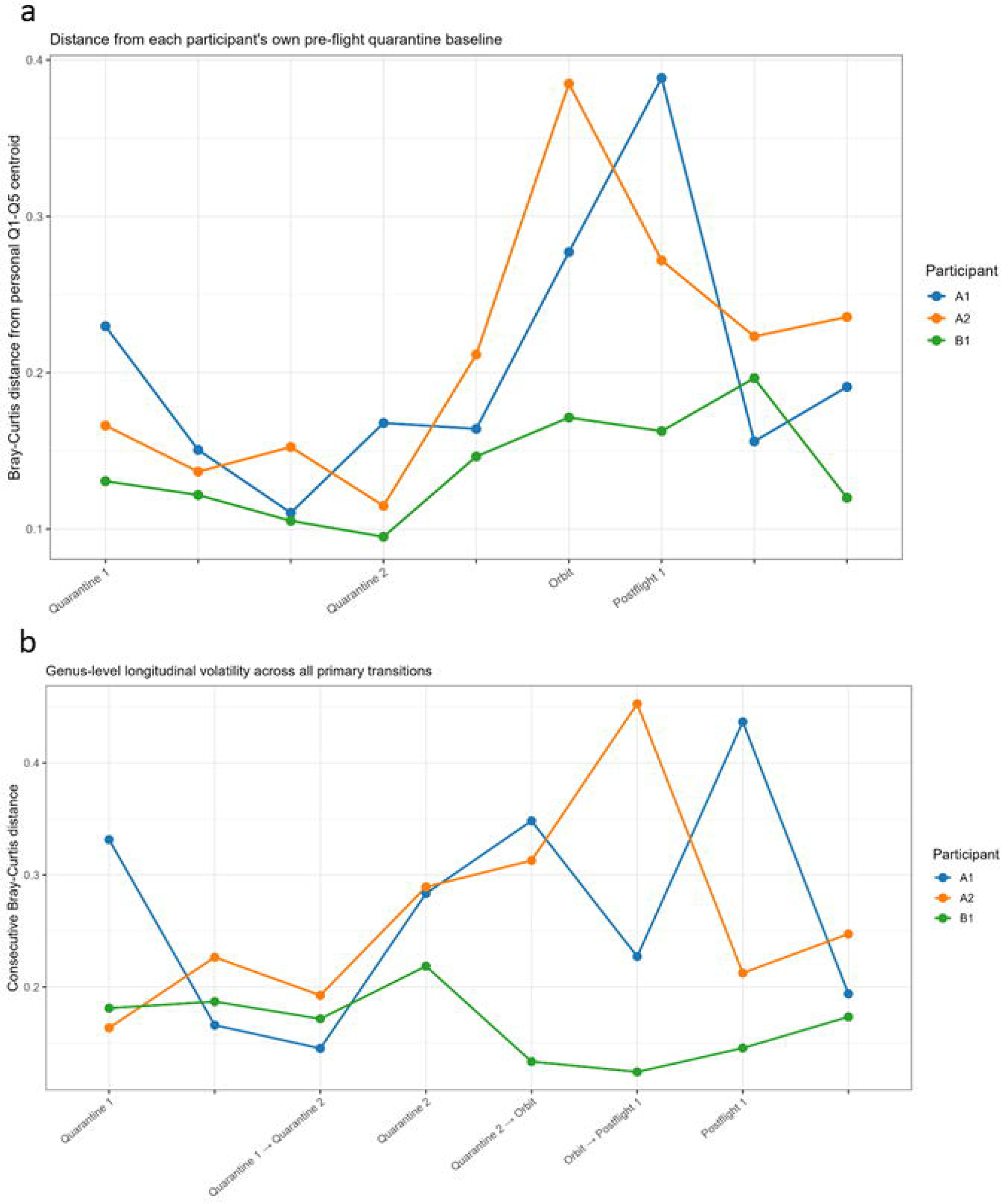
Personal-baseline displacement and longitudinal volatility. a, Bray-Curtis distance of each sample from the centroid of that participant’s Q1-Q5 genus-level profiles. **b,** Bray-Curtis distance between consecutive phase-matched windows. Lines display individual participants; astronaut-versus ground-based participant summaries are descriptive and were not treated as independent group-level hypothesis tests. Because sampling intervals differed, a time-gap-adjusted sensitivity analysis is provided in Supplementary Fig. 3.

Consecutive Bray-Curtis distances were calculated across all eight transitions from Q1 to PostF3. Mean step distance was 0.264 in A1 and A2 combined and 0.167 in B1, a 1.58-fold difference (Fig. 4b). The largest astronaut-to-B1 ratios occurred from Q5 to O1 (2.48-fold), O1 to PostF1 (2.74-fold) and PostF1 to PostF2 (2.23-fold). The timing was participant-specific: A2 showed its largest step from O1 to PostF1, whereas A1 showed its largest step from PostF1 to PostF2. Transition-level values and the time-gap-adjusted sensitivity plot are provided in Supplementary Table 6 and Supplementary Figure 3.

### Taxon-level changes were selective, with a significant quarantine-to-orbit increase in Collinsella

Participant-baseline-adjusted CLR analysis of the full A1/A2/B1 dataset identified genera with large O1 astronaut-to-ground-based-participant contrasts (Fig. 5a). *Wujia*, *Streptococcus*, *Odoribacter*, *Agathobacter*, *Paraprevotella*, *Fastidiosipila* and *Lachnoclostridium* decreased relative to personal baseline in both astronauts and more strongly than in B1. *Hominenteromicrobium*, *Caproiciproducens*, *Acutalibacter*, *Ruthenibacterium*, *Denitrobacterium*, *Raoultibacter* and *Collinsella* increased in both astronauts. *Kineothrix* had the largest mean contrast, but A1 and A2 changed in opposite directions and it was therefore not interpreted as a shared astronaut pattern. Participant-specific genus-level compositional trajectories across the four primary mission phases are shown in Fig. 6.

**Figure 5.**
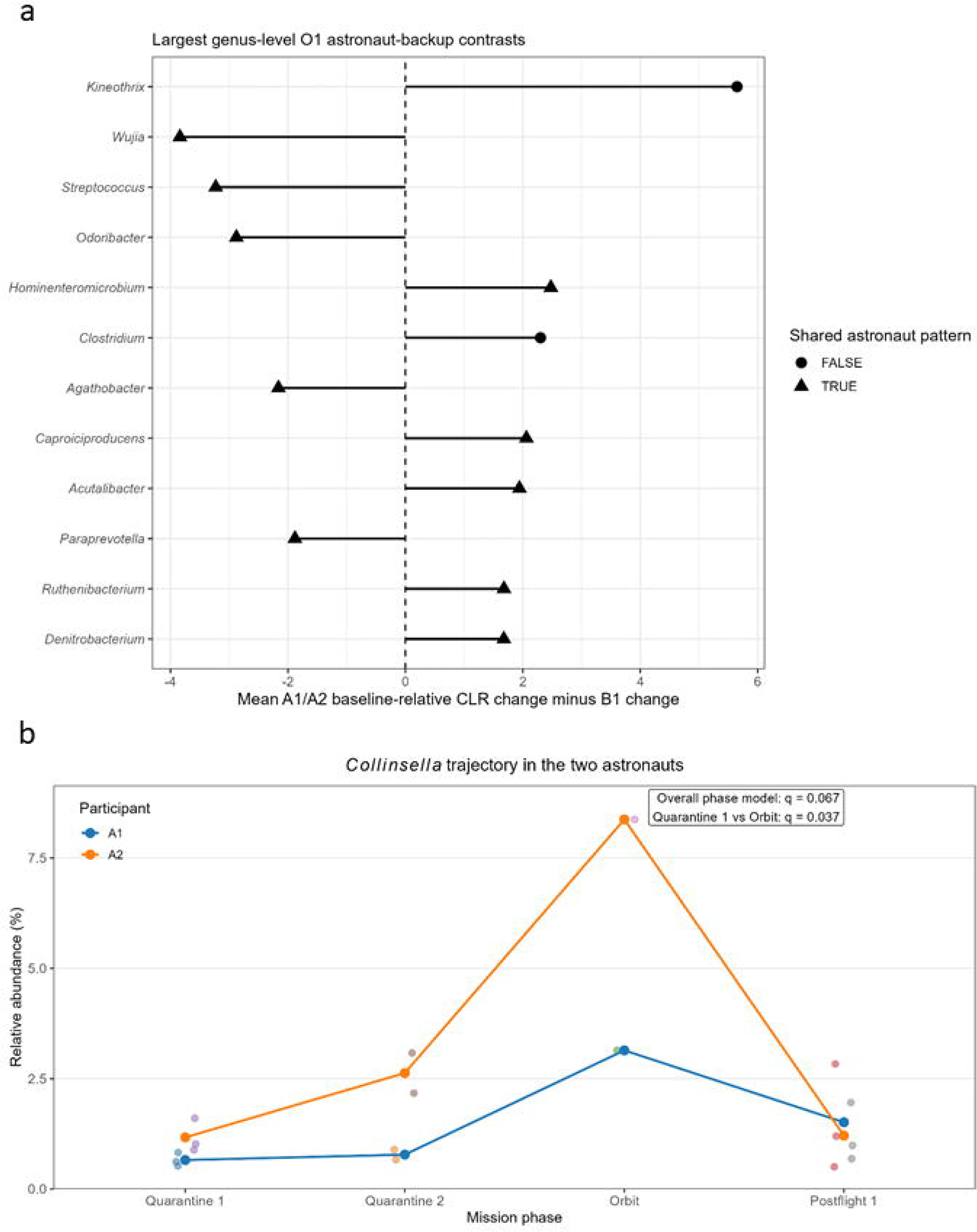
Baseline-adjusted and astronaut-only genus patterns. a, Largest O1 astronaut-to-ground-based-participant contrasts calculated as the mean A1/A2 baseline-relative CLR change minus the corresponding B1 change. Triangles indicate genera that changed in the same direction in both astronauts and for which each astronaut’s absolute change exceeded the B1 change. **b,** *Collinsella* relative abundance in A1 and A2 across the four primary mission phases. Small points show individual samples and connected points show participant-specific phase means. *Collinsella* was associated with mission phase in the astronaut-only CLR mixed model (q = 0.067) and increased from Quarantine 1 to Orbit in MaAsLin2 (coefficient = 2.586, q = 0.037).

**Figure 6.**
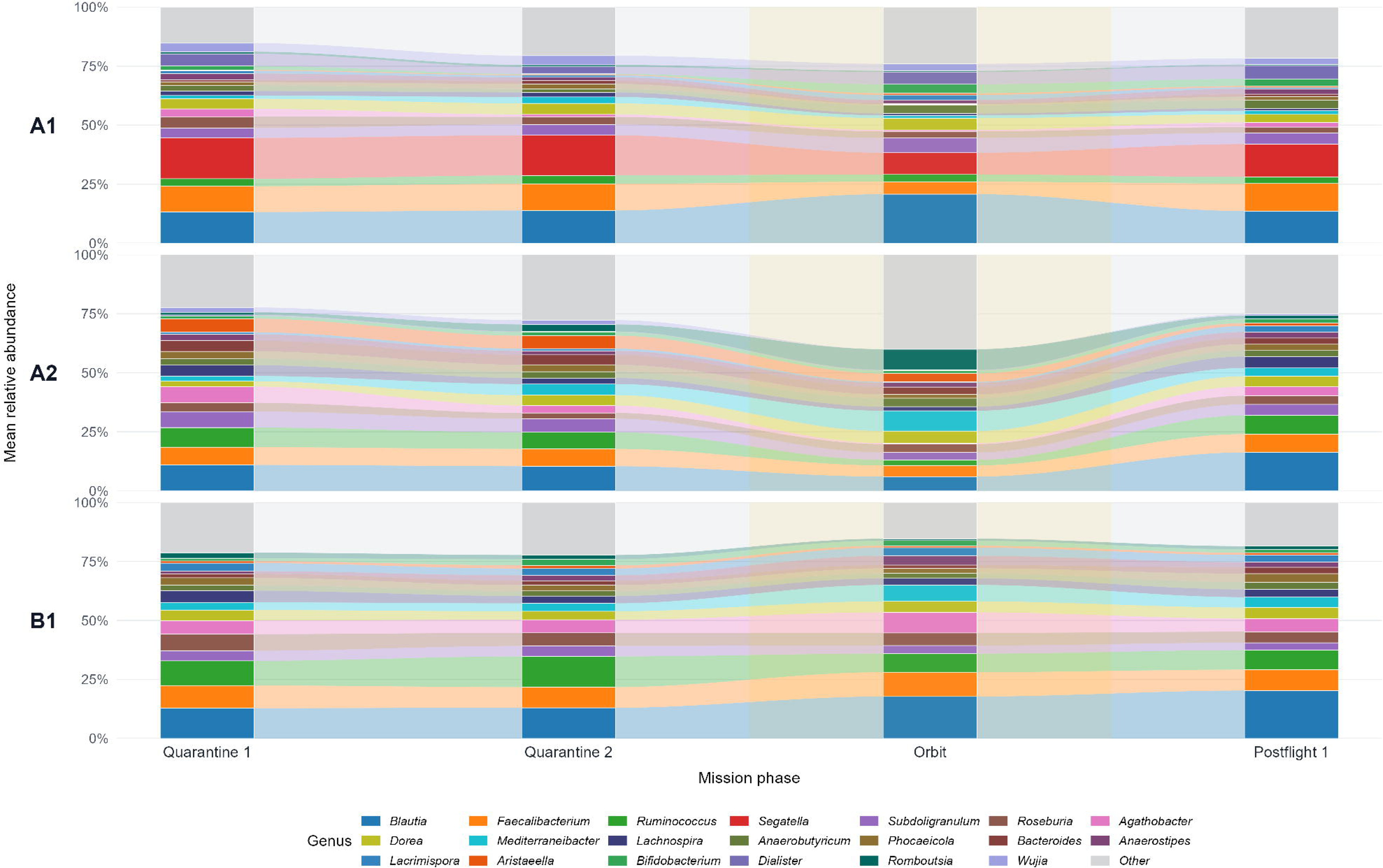
Genus-level compositional trajectories across mission phases. Stacked alluvial plots show mean genus-level relative abundances for astronauts A1 and A2 and the ground-based participant B1 across Quarantine 1, Quarantine 2, Orbit and Postflight 1. Colored bands represent individual genera and connect phase-specific mean abundances to visualize longitudinal changes in community composition within each participant. The 20 genera with the highest mean relative abundance across the dataset are displayed individually; all remaining genera are grouped as Other.

We next treated the astronaut-only mixed models as the primary taxon-level phase analysis, thereby testing systematic changes across A1 and A2 without requiring the ground-based participant to follow the same trajectory. Using the prespecified criteria q ≤ 0.10 and fixed-effect variance fraction ≥ 0.20, 2 of 81 genera and 5 of 139 species were phase-associated across the four primary phases. The associated genera were *Odoribacter* (p = 0.00076, q = 0.0617; fixed-effect variance fraction = 0.621) and *Collinsella* (p = 0.00166, q = 0.0673; fixed-effect variance fraction = 0.584). The associated species were *Bacteroides uniformis* (q = 0.0721), *Odoribacter splanchnicus* (q = 0.0882), *Bacteroides thetaiotaomicron* (q = 0.0882), *Bifidobacterium pseudocatenulatum* (q = 0.0882) and *Collinsella aerofaciens* (q = 0.0948).

Mission phase was the dominant modeled variance component for the first four species, but not for Collinsella *aerofaciens*. In the focused Quarantine 1-versus-Orbit MaAsLin2 comparison, *Collinsella* increased with a coefficient of 2.586 (p = 0.00045, q = 0.0365), while *Novisyntrophococcus* (q = 0.0605) and *Acutalibacter* (q = 0.0794) also passed q ≤ 0.10. *Collinsella* increased at O1 to 3.14% in A1 and 8.37% in A2, followed by participant-specific recovery (Fig. 5b). Astronaut-only taxon counts and associated taxa are listed in Supplementary Tables 7 and 8.

As a sensitivity analysis, the full three-participant fixed-effect models asked whether a phase effect was shared across A1, A2 and B1. These models remained dominated by participant identity: using q ≤ 0.10 and partial R2 ≥ 0.20, 68 of 86 genera and 128 of 151 species were participant-associated, whereas no genus or species met the mission-phase criterion after rank-wise correction. *Eubacteriales* was the only corrected order-level phase-associated feature. This sensitivity result does not negate the astronaut-only associations; it indicates that they were not universal trajectories shared by the two astronauts and the ground-based participant (Supplementary Table 9).

### Predicted KEGG functional pathway analysis

Differential analysis of predicted KEGG ortholog abundances (KOs) was first performed using DESeq2 for each primary mission phase relative to Quarantine 1. One single significantly differentially abundant KO was identified in the Quarantine 2 vs. Quarantine 1 and Postflight 1 vs. Quarantine 1 comparison, whereas 116 predicted KOs showed significant differential abundance in the Orbit vs. Quarantine 1 comparison. For each contrast, the complete tested KO set was subsequently ranked by shrunken log2 fold change and used as input for GSEA. Although 11 pathways met the statistical GSEA threshold, their leading-edge KOs showed very small effect sizes; we therefore report these results in Supplementary Table 10 and refrain from biological interpretation.

In the Orbit vs. Quarantine 1 comparison, GSEA identified three significantly enriched pathways, with Galactose metabolism (NES = 1.65, FDR = 0.042) and Aminoacyl-tRNA biosynthesis (NES = 1.68, FDR = 0.0089) positively enriched in Orbit, whereas the Two-component system pathway was negatively enriched (NES = −1.46, FDR = 0.044). The leading-edge subsets comprised 7, 3, and 41 core KOs for the Galactose metabolism, Aminoacyl-tRNA biosynthesis, and Two-component system pathways, respectively (Fig. 7). Consistent with the GSEA results, ORA identified Galactose metabolism as the only significantly overrepresented pathway (Rich Factor = 0.140, 6 KOs, p = 4.34 × 10⁻, adjusted p = 0.0208).

**Figure 7.**
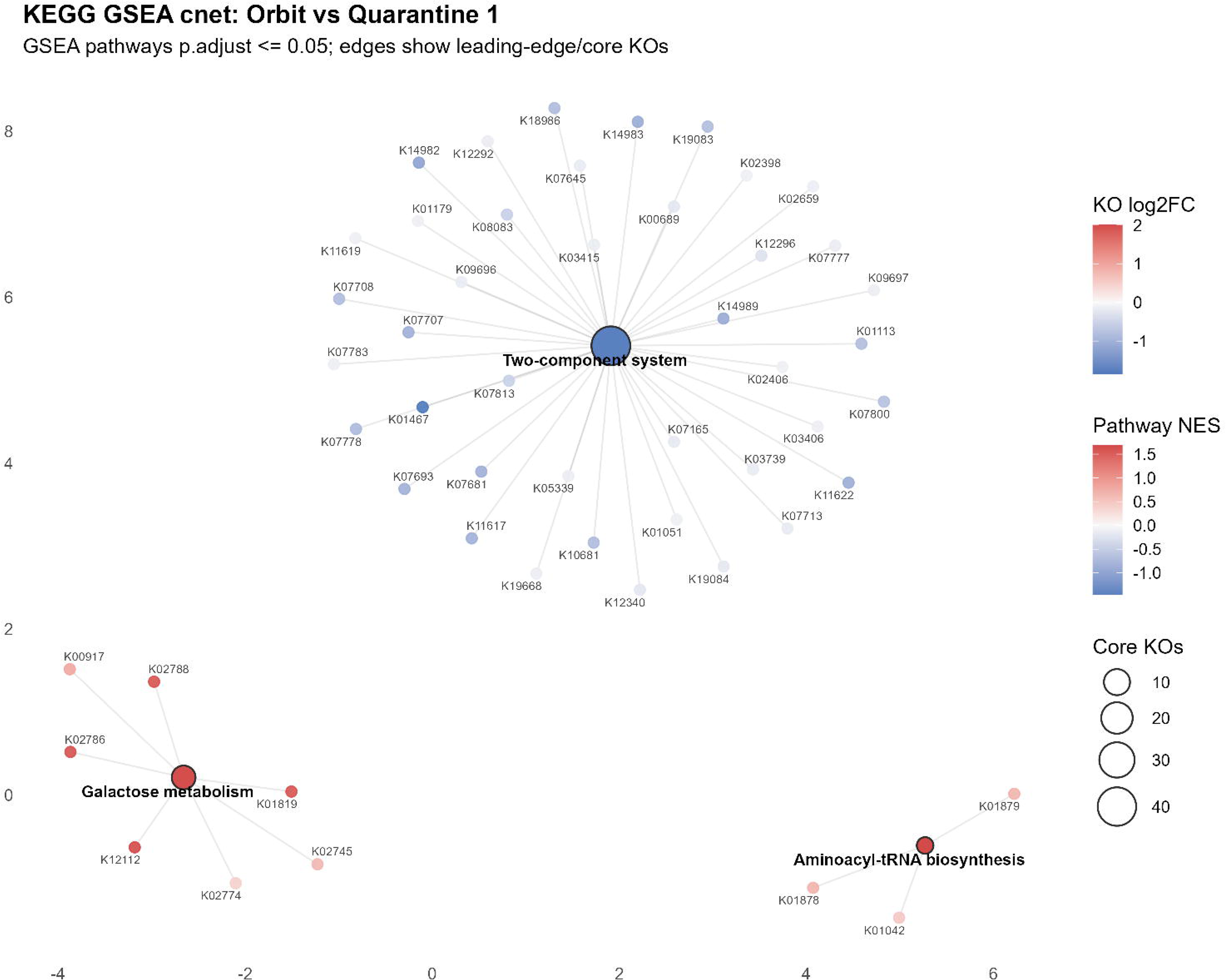
KEGG pathway enrichment network for the Orbit vs. Quarantine 1 comparison. Significantly enriched pathways identified by GSEA are shown as large nodes connected to their leading-edge (core) KEGG orthologs (KOs). Pathway node color represents the normalized enrichment score (NES), with positive enrichment shown in red and negative enrichment in blue. KO node color indicates the corresponding log2 fold change, and pathway node size is proportional to the number of leading-edge KOs contributing to pathway enrichment.

## Discussion

This study provides a longitudinal full-length 16S view of fecal microbiota dynamics during a short-duration International Space Station mission. The central result is not a uniform restructuring of the gut community, but a combination of persistent personal signatures and temporally localized changes. Across A1, A2 and B1, participant identity explained substantially more beta-diversity variation than mission phase. When the community-phase question was restricted to the two flown participants, however, mission phase accounted for 23.4% of genus-level and 22.3% of species-level Bray-Curtis variation. This distinction is important: global community structure remained individualized, while the astronaut-only analysis supported a flight-window-associated community signal.

Distance from personal Q1-Q5 centroids and consecutive-window volatility provided the clearest indication of increased microbiome instability around the flight and early-recovery windows. Both measures increased around O1 and early recovery in the flown participants, although the timing differed between A1 and A2. The ground-based participant provides useful calendar-matched context, but cannot control for every environmental, dietary or behavioral exposure. The taxon-level results likewise support selective rather than community-wide change. Several genera moved in the same baseline-adjusted direction in both astronauts at O1. In the primary astronaut-only phase screen, 2 genera and 5 species met the prespecified exploratory criteria. *Odoribacter* and *Collinsella* were phase-associated at genus-level, while the species-level set comprised *Bacteroides uniformis*, *Odoribacter splanchnicus*, *Bacteroides thetaiotaomicron*, *Bifidobacterium pseudocatenulatum* and *Collinsella aerofaciens*. *Collinsella* showed the strongest genus-level evidence of a flight-window-associated change: it passed FDR correction in the focused Quarantine 1-versus-Orbit comparison and also met the q ≤ 0.10 exploratory criterion in the overall astronaut-only phase model. The absence of corrected genus- or species-level phase effects in the A1/A2/B1 sensitivity model shows that these patterns were not universal across all three participants, rather than invalidating the astronaut-only findings. These associations provide additional support for selective flight-window-associated responses, but they remain observations in two astronauts and are not population-level biomarkers or evidence that microgravity alone caused the changes.

The findings are consistent with previous astronaut studies in which microbiome changes occurred against strong individual-specific backgrounds and varied in magnitude and persistence. Recent short-duration multi-omics investigations similarly identified measurable but heterogeneous host and microbial responses. Controlled models also support the biological plausibility of space-relevant exposures influencing host-microbiota interactions, although the present observational design cannot attribute changes specifically to microgravity.^9,10,11,12,19,20^

Functional prediction further suggested positive enrichment of predicted galactose metabolism and aminoacyl-tRNA biosynthesis pathways and negative enrichment of the predicted two-component system pathway during the flight-associated window. Notably, increased predicted galactose metabolism was supported by both GSEA and ORA in our dataset and is consistent with findings from space-flown mice, in which shotgun metagenomic analysis independently identified over-representation of the same KEGG galactose metabolism pathway during spaceflight.^19^ Spaceflight-associated alterations in inferred microbial carbohydrate utilization have also been reported in an independent ISS mouse study.^21^ Aminoacyl-tRNA biosynthesis has likewise been implicated in intestinal metabolic responses to simulated weightlessness in rats, although it was decreased in that metabolomic study, whereas our marker-gene-based prediction indicated an increase.^22^ This difference may reflect differences in host species, experimental conditions, biological layer measured, or the distinction between direct metabolomic measurements and predicted microbial functional potential. The decrease in predicted two-component system capacity may represent a more study-specific response. These functional results should nevertheless be interpreted as marker-gene-based predictions rather than direct measurements of microbial gene content or activity.

The separation of primary early-recovery samples from later travel-associated samples was deliberately conservative. PostF4-PostF6 may contain biologically meaningful information about real-world recovery, but those windows also incorporate changes in geography, diet, circadian timing and environmental exposure. They should therefore be retained as a supplementary sensitivity analysis rather than combined with the controlled primary inference.

Several limitations constrain interpretation. The cohort comprised only two astronauts and one ground-based participant, the in-flight fecal sampling density was limited, and the study was observational. The astronaut-only community and taxon models provide useful longitudinal assessments of phase-associated patterns in the flown participants, but population-level estimates cannot be derived from two astronauts. Consecutive distances and astronaut means are descriptive, not independent replicates. In addition, the astronauts underwent a pre-flight bowel preparation procedure before the late-quarantine sampling windows, which may have contributed to microbiota changes preceding the in-flight sample. Full-length 16S sequencing improves marker-gene resolution but does not provide direct functional measurements and may not resolve all species accurately. Finally, spaceflight integrates microgravity, radiation, confinement, diet, stress, sleep disruption and travel, whose individual contributions cannot be disentangled in this design.

In summary, short-duration spaceflight was accompanied by an astronaut-only community-phase signal, increased displacement from personal microbiota baselines and higher longitudinal volatility in the flown participants, while global community structure remained strongly individualized. The primary astronaut- only taxon models identified 2 associated genera and 5 associated species, including a significant quarantine-to-orbit increase in *Collinsella*. Longitudinal full-length 16S profiling may therefore be useful for defining personal microbial baselines and identifying temporally localized community and taxon-level responses in sample-limited astronaut studies.

## Methods

### Ethics approval and informed consent

All procedures involving human participants were conducted under the “Mapping Astronaut Meta- GenOmics: a Microbial Profiling Research” (MAGOR) protocol. The study was approved by the NASA Institutional Review Board (eIRB STUDY00000787; 6 March 2025), the Human Research Multilateral Review Board (STUDY787; 23 April 2025), and the ESA Medical Board. Preliminary approval was provided by the Hungarian Medical Research Council on 25 November 2024, contingent on NASA approval. The study was conducted in accordance with the Declaration of Helsinki and applicable institutional, national, and international regulations. Written informed consent was obtained from all participants before enrolment, and all data were pseudonymized.

Study participants and sample collection

The study included three male volunteers aged 30-45 years: two astronauts who participated in the International Space Station mission and one ground-based participant who remained on Earth but followed the same daily schedule. Samples were collected using DNA/RNA Shield Fecal Collection Tubes (Zymo Research, R1101-E). Samples collected on Earth and aboard the International Space Station were stored at -80 °C and remained frozen during storage and transport.

The complete sequencing dataset contained 47 samples, of which 27 phase-matched fecal samples from A1, A2 and B1 constituted the primary analysis dataset (Supplementary Table 1). Samples were assigned to phase-matched windows according to collection day relative to launch. The primary analysis comprised Q1-Q5, O1 and PostF1-PostF3. PostF4-PostF6 were excluded from primary inference because they coincided with substantial post-mission travel and were reserved for sensitivity analysis.

### DNA extraction

Genomic DNA was isolated from 800 µl of DNA/RNA Shield-preserved fecal material using the ZymoBIOMICS 96 MagBead DNA Kit (Zymo Research, D4308-E). Samples were lysed in bead-containing tubes using the kit lysis solution. Clarified lysates were combined with MagBinding Buffer and magnetic beads, washed sequentially with kit binding and wash buffers, and eluted in 50 µl ZymoBIOMICS DNase/RNase-Free Water. DNA was stored at -80 °C. Yield was measured by Qubit fluorometry and integrity was evaluated using an Agilent TapeStation.

### Full-length 16S amplification library preparation and sequencing

The bacterial V1-V9 16S rRNA region was amplified using modified Oxford Nanopore primers 27F (5′- AGRGTTYGATYMTGGCTCAG-3′) and 1492R (5′-CGGYTACCTTGTTACGACTT-3′). PCR comprised 95 °C for 1 min; 25 cycles of 95 °C for 20 s, 55 °C for 30 s and 65 °C for 2 min; and 1 cycle of 65 °C for 5 min. Amplicons were purified using AMPure XP beads, washed with 70% ethanol and eluted in 10 µl nuclease-free water.23

Libraries were prepared using the Oxford Nanopore Ligation Sequencing Amplicons workflow and Native Barcoding Kit 96 V14 (SQK-NBD114.96). Concentration and fragment-size distribution were evaluated using a Qubit 4 Fluorometer and Agilent 4150 TapeStation. Barcoded libraries were normalized, pooled and sequenced on a PromethION platform. Basecalling was performed with Dorado v7.11.2 in super-accurate mode.

### Sequence processing and Kraken2 classification

Demultiplexed full-length 16S FASTQ files were processed using Filtlong (v0.3.1) with the following parameters: --min_length 1500 --max_length 1800. Only reads between 1,500 and 1,800 bp were retained for downstream analyses.

The primary taxonomy used in this manuscript was generated by the Kraken2 analysis branch. Full-length reads were classified against a Kraken2 standard database,^24^ and classification-derived count and taxonomy tables were imported as a phyloseq object.^25^

Kraken2 version and database metadata: Kraken2 v2.1.7 with the k2_standard_20260226 database.

Demultiplexed full-length 16S FASTQ files were processed using NanoASV with a minimum read quality score of 8, minimum ASV abundance of 5, a maximum subsample of 2,000,000 reads per barcode and 40 threads. NanoASV used the SILVA 138.2 SSURef database for its internal taxonomic processing.^26,27^

### Sample selection and taxon filtering

Sample metadata were curated in R and merged with the Kraken2 phyloseq object. Samples outside the matched longitudinal framework, reserve or non-target samples, samples with missing time-period information and samples with fewer than 1,000 reads were excluded. At species-level, taxa were first required to reach at least 10 reads and a relative abundance of 1 × 10^-5 in at least one sample. The retained Kraken2 rows were then required to be present in at least three samples, have at least 1,000 total reads, reach at least 100 reads in one sample and reach a maximum within-sample relative abundance of 0.001. All four second-stage criteria had to be satisfied. This yielded 152 retained Kraken2 rows, resolving to 5 phyla, 16 orders, 86 genera and 151 named species in the primary analysis.

Mission-phase groups were Quarantine 1 (Q1-Q3), representing the pre-flight quarantine period, Quarantine 2 (Q4-Q5), following the NASA pre-flight bowel preparation procedure for astronauts A1 and A2, Orbit (O1), and Postflight 1 (PostF1-PostF3). The late Postflight 2 group (PostF4-PostF6) was excluded from the primary analysis. Relative-abundance figures used within-sample total-sum normalization.

### Community composition ordination and multivariate testing

Taxa were agglomerated independently at phylum, order, genus and species-levels. Within-sample relative abundances were used to calculate Bray-Curtis dissimilarities. Genus-level non-metric multidimensional scaling used vegan metaMDS with two dimensions, up to 200 random starts, no additional transformation and a fixed random seed of 123. An Aitchison sensitivity analysis used a pseudocount of 0.5, centered log-ratio transformation and principal-component analysis.

Marginal PERMANOVA included participant identity and mission phase in the same model and used adonis2 with by = "margin" and 9,999 permutations. The 27-sample A1/A2/B1 model was used to quantify overall participant structure and to contextualize the ground-based participant. The astronaut- specific community-phase test was then performed on the 18 A1/A2 samples using the same rank- agglomerated Bray-Curtis framework. Homogeneity of multivariate dispersion was evaluated from Bray-Curtis PCoA distances to mission-phase spatial medians with 9,999 label permutations.

### Personal-baseline displacement and longitudinal volatility

For each participant, the personal preflight baseline was defined as the centroid of Q1-Q5 genus-level relative-abundance profiles. Bray-Curtis distance from this centroid was calculated for every retained window. Aitchison distance from the corresponding CLR centroid was retained as a sensitivity measure.

Longitudinal volatility was calculated as Bray-Curtis distance between each pair of consecutive windows: Q1-Q2, Q2-Q3, Q3-Q4, Q4-Q5, Q5-O1, O1-PostF1, PostF1-PostF2 and PostF2-PostF3. Participant-specific distances and the arithmetic mean of A1 and A2 were reported descriptively. No formal astronaut-to-ground-based participant hypothesis test was performed because the study contained two flown participants and one ground-based participant.

### Baseline-adjusted taxon trajectories and exploratory models

Genus count matrices were transformed using a pseudocount of 0.5 and the centered log-ratio transformation. For each participant and genus, baseline-relative CLR change was calculated by subtracting the mean Q1-Q5 CLR abundance. At O1, the astronaut-to-ground-based-participant contrast was defined as the mean A1/A2 baseline-relative change minus the B1 change. A robust descriptive candidate required A1 and A2 to change in the same direction and each astronaut’s absolute change to exceed the B1 change.

Two complementary taxon-modeling strategies were used. The primary phase-associated taxon screen was restricted to the two astronauts. At each taxonomic rank, CLR-normalized abundance was modeled with mission phase as a fixed effect and participant as a random intercept. Taxa were classified as phase-associated when the rank-wise Benjamini-Hochberg-adjusted q value was <= 0.10 and the fixed-effect variance fraction was ≥ 0.20; phase was classified as dominant when its variance fraction exceeded the participant fraction. A focused MaAsLin2 model compared Quarantine 1 with Orbit in A1 and A2. As a sensitivity analysis, the full A1/A2/B1 dataset was modeled using CLR abundance as the response and participant identity plus mission phase as fixed effects; partial F tests compared the full model with models omitting participant or phase. Benjamini-Hochberg correction was applied separately within each taxonomic rank or focused comparison. The prespecified exploratory threshold was q ≤ 0.10; results with q < 0.05 were considered significant.^28^ We intentionally used q ≤ 0.10 as an exploratory FDR threshold because statistical power was limited by the small number of samples, especially the single in-flight/orbit sampling window per participant. To reduce the risk of overinterpreting weak signals, we combined this threshold with a fixed-effect variance fraction criterion of ≥ 0.20. We therefore do not present these taxa as conventionally significant at q <= 0.05, but as biologically plausible, effect- supported candidates relevant to short-duration spaceflight-associated microbiota dynamics.

### Functional prediction and KEGG enrichment analysis

Functional prediction was performed as an exploratory complement to the primary Kraken2 taxonomic analysis. The NanoASV/PICRUSt2 branch used NanoASV ASV abundance tables from the same full-length 16S reads to infer unstratified KEGG ortholog (KO) metagenome abundances.^29,30^ The exported PICRUSt2 KO table was converted to a phyloseq object and merged with the curated MAGOR metadata. The manuscript-facing functional analysis used the same phase windowing as the strict primary analysis, retained Q1-Q5, O1 and PostF1-PostF3, excluded PostF4-PostF6, and was restricted to the two flown participants.

Samples with fewer than 1,000 total predicted KO counts were removed before downstream testing. Low-support KOs were filtered before ordination and differential analysis using a permissive support rule: KOs were retained if they were present in at least three samples, had a total predicted count of at least 100, had a maximum single-sample count of at least 25, or reached a maximum within-sample relative abundance of at least 1 x 10^-6.

Predicted KO composition was assessed by Hellinger transformation followed by Euclidean-distance marginal PERMANOVA with participant identity and mission phase in the same model (adonis2, by = "margin", 9,999 permutations). DESeq2 used a design including participant identity and mission phase (∼ participant + phase), with Quarantine 1 as the reference phase.^31^ DESeq2 was run with the Wald test, local dispersion fitting, positive-count size-factor estimation, independent filtering and Cook’s-distance outlier replacement/filtering disabled. Log2 fold changes were shrunk using adaptive shrinkage.^32^ Contrasts compared Quarantine 2, Orbit and Postflight 1 with Quarantine 1; Postflight 2 was excluded upstream from the primary inference. KOs were considered differentially abundant at Benjamini-Hochberg adjusted padj <= 0.05, with no additional absolute log2 fold-change cutoff.

KEGG enrichment analyses were performed with clusterProfiler.^33^ Over-representation analysis used enrichKEGG for KEGG pathways and enrichMKEGG for KEGG modules, with significant KOs as the foreground set and all tested KOs as the universe. Gene-set enrichment analysis used the complete KO list ranked by shrunken log2 fold change and gseKEGG with organism = "ko" and keyType = "kegg". Pathways, modules and GSEA terms were retained at p.adjust <= 0.05. ORA and GSEA were interpreted separately: ORA tests whether thresholded significant KOs are over-represented in a pathway or module, whereas GSEA tests whether all KOs in a pathway show a coordinated shift in rank even when few individual KOs pass the DESeq2 threshold. Dotplots, heatmaps, cnet-style networks, chord diagrams and KEGG map overlays were generated for visual review; KEGG map overlays used Pathview.^34^ Because PICRUSt2 provides marker-gene-based functional predictions, the functional results were interpreted as predicted pathway capacity rather than direct metagenomic, metatranscriptomic or metabolomic measurements.

### Statistics and reproducibility

The primary community dataset contained 27 samples from three participants, while astronaut-only community and taxon analyses contained 18 samples from A1 and A2; the Quarantine 1-versus-Orbit comparison contained eight samples. Exact n values are reported in figure legends and Table 1. All multivariate tests were two-sided permutation tests. Actual p values are reported where available; p values below 0.001 are reported as p < 0.001 in the narrative. The alpha level was 0.05 for community-level tests. Taxon-level analyses used Benjamini-Hochberg correction and the prespecified q ≤ 0.10 exploratory threshold, with q < 0.05 identified explicitly. No data points were excluded on the basis of the observed outcome. Exported source result tables supporting the article-specific analyses are included in the generated submission package, and manuscript-facing statistics are audit-checked against these files.

## Acknowledgements

The authors thank Dr Balázs Nagy, Dr Boldizsár Balázs, István Örökös-Tóth and the HUNOR Hungarian Astronaut Program team for administrative, logistical and technical support. We thank Axiom Space for mission coordination and support with the NASA ethics process, ESA for sample-collection logistics in Europe and the mission operations teams involved in study implementation.

This project was supported by the HUNOR Hungarian to Orbit Program of the Ministry of Foreign Affairs and Trade to DT, the Hungarian Academy of Sciences Momentum Grant LA1020-8/2020 to DT and National Research Development and Innovation Office grants FK 142676 to DT and K 142674 and ADV 152705 to ZB. The funders had no role in study design, data collection, analysis, interpretation or manuscript preparation. The article-processing charge was supported by the University of Szeged Open Access Fund grant 8855 to DT.

## Author contributions

DT conceived and supervised the study. DT, GG and IP led project administration and coordination with the participating spaceflight organizations, with additional contributions from ZC and ZB. ÁD performed the laboratory work and sequencing-related sample processing. GG and DT performed the initial data analyses and interpretation. BK performed and integrated the primary statistical and bioinformatic analyses underlying the final manuscript. DT contributed to the design of data visualizations, and IP prepared the study-design figure. TJ contributed to bioinformatic analysis. ÁTS contributed clinical and mission-related expertise. GG, BK, DT and ZB interpreted the results. GG and DT prepared the original manuscript draft, and BK and ZB made substantial contributions to subsequent writing and revision. All authors reviewed and approved the final manuscript.

## Competing interests

The authors declare no financial or non-financial competing interests.

## Notes

### Competing Interest Statement

The authors have declared no competing interest.

