## Supplementary figures and images for "Full-length 16S profiling reveals individualized gut microbiota dynamics during short-duration spaceflight"

### Supplementary Figure 1

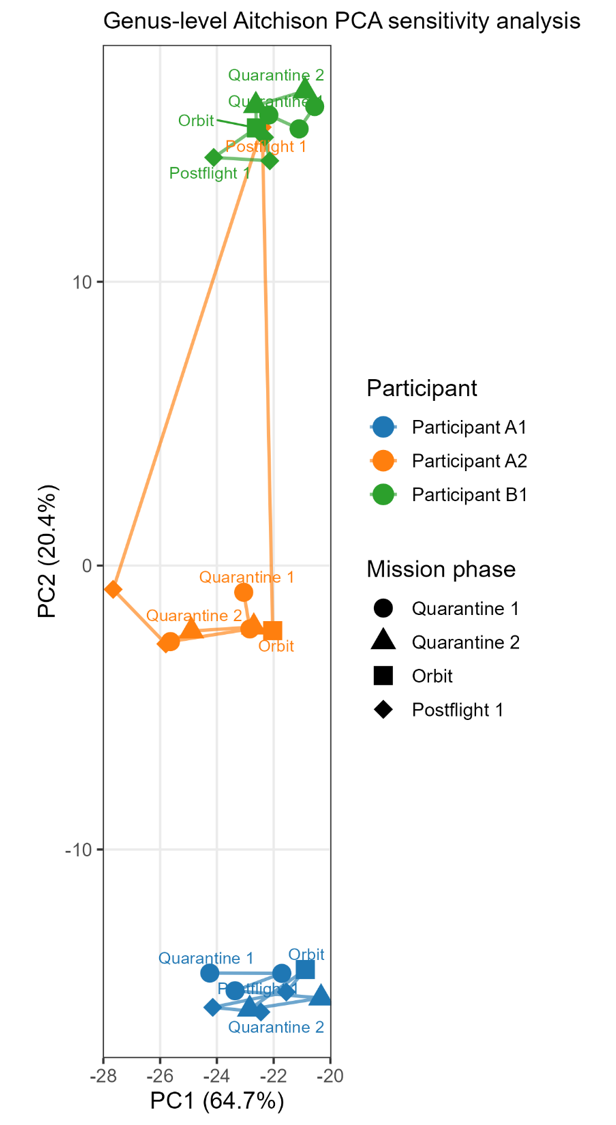

### Supplementary Figure 2

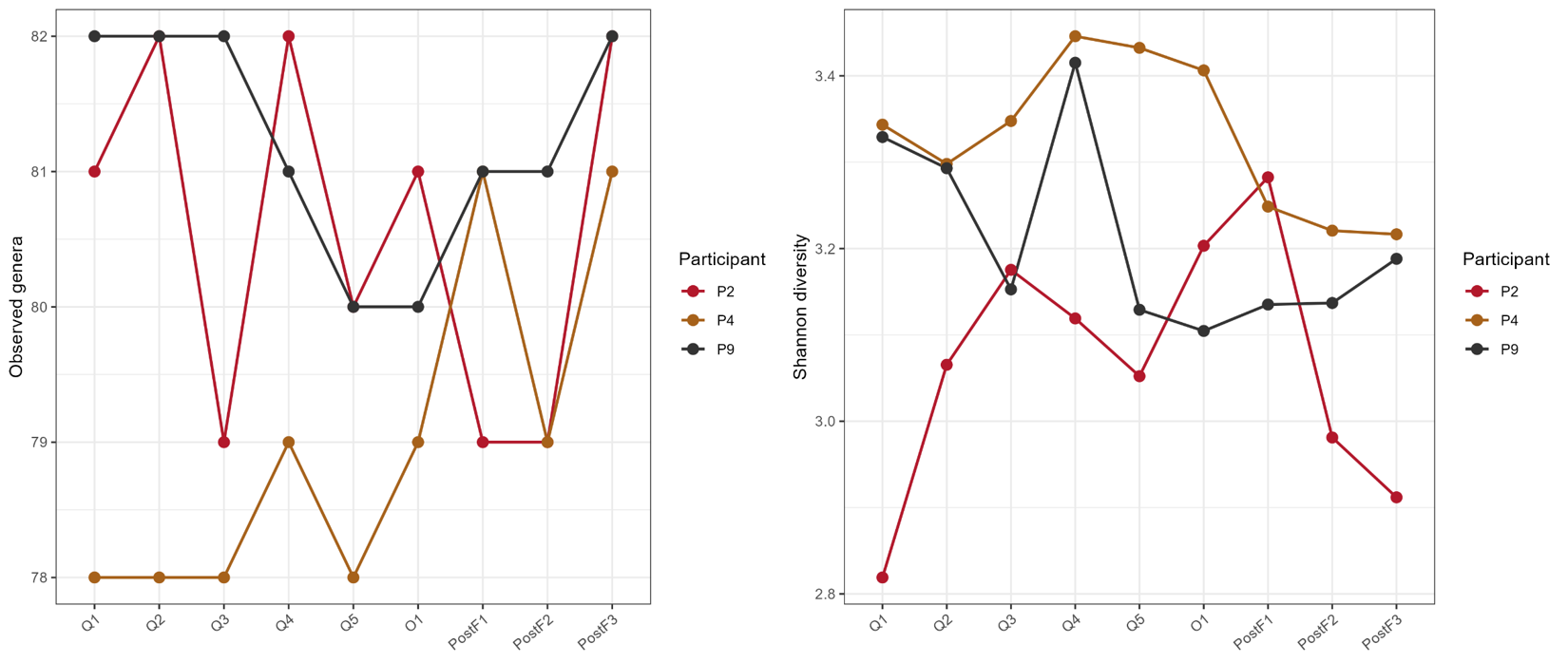

### Supplementary Figure 3

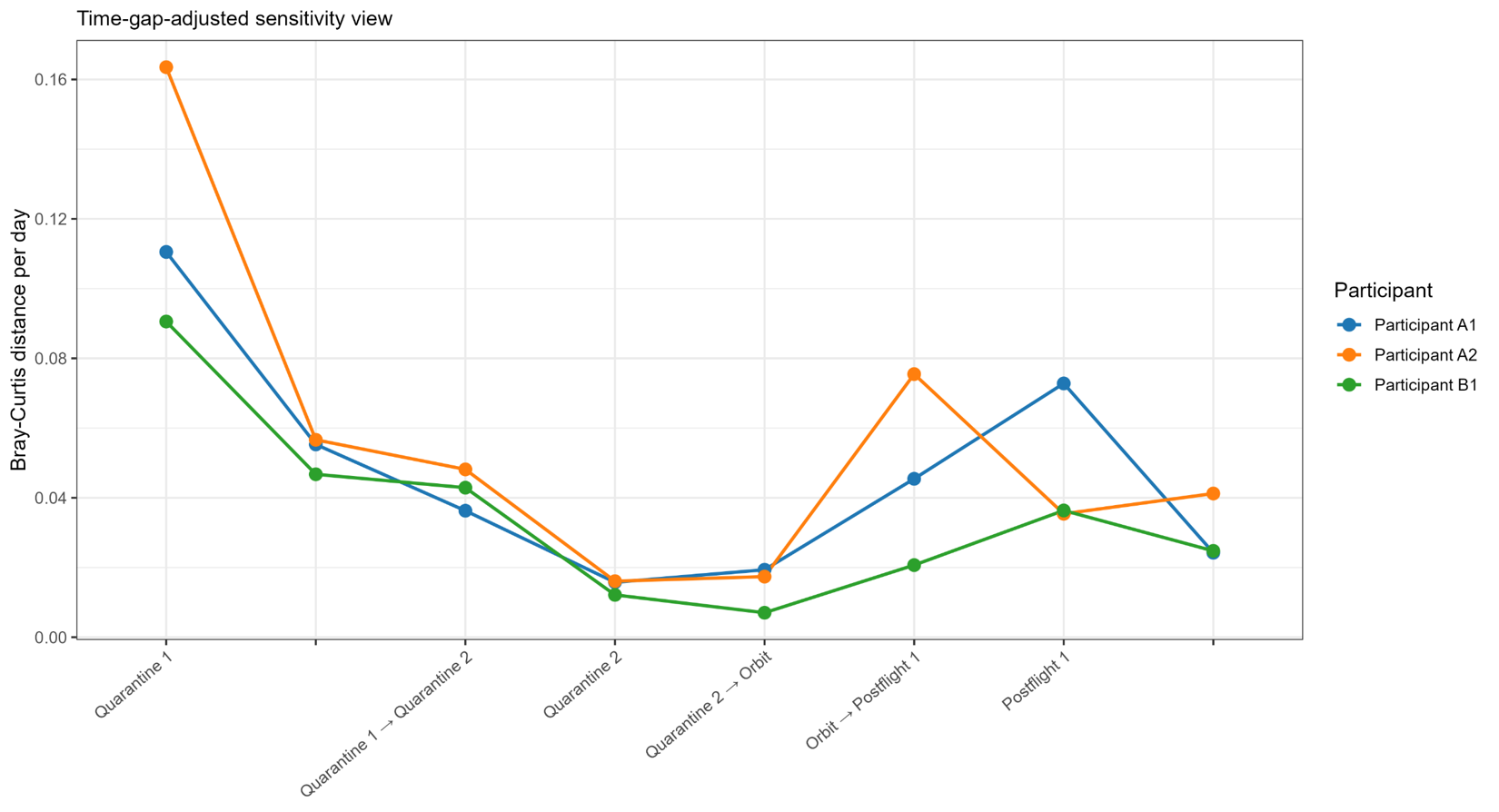
